# Representational organization of novel task sets during proactive encoding

**DOI:** 10.1101/710244

**Authors:** Ana F. Palenciano, Carlos González-García, Juan E. Arco, Luiz Pessoa, María Ruz

**Affiliations:** Mind, Brain, and Behavior Research Center (CIMCYC), University of Granada, 18011, Granada, Spain.; Department of Experimental Psychology, Ghent University, 9000, Ghent, Belgium.; Psychology Department, University of Maryland, 20742, Maryland, United States of America

**Author notes:** Corresponding author: María Ruz. E-mail address (M. Ruz).

## Abstract

Recent multivariate analyses of brain data have boosted our understanding of the organizational principles that shape neural coding. However, most of this progress has focused on perceptual visual regions (Connolly et al., 2012), whereas far less is known about the organization of more abstract, action-oriented representations. In this study, we focused on humans’ remarkable ability to turn novel instructions into actions. While previous research shows that instruction encoding is tightly linked to proactive activations in fronto-parietal brain regions, little is known about the structure that orchestrates such anticipatory representation. We collected fMRI data while participants (both males and females) followed novel complex verbal rules that varied across control-related variables (integrating within/across stimuli dimensions, response complexity, target category) and reward expectations. Using Representational Similarity Analysis (Kriegeskorte et al., 2008) we explored where in the brain these variables explained the organization of novel task encoding, and whether motivation modulated these representational spaces. Instruction representations in the lateral prefrontal cortex were structured by the three control-related variables, while intraparietal sulcus encoded response complexity and the fusiform gyrus and precuneus organized its activity according to the relevant stimulus category. Reward exerted a general effect, increasing the representational similarity among different instructions, which was robustly correlated with behavioral improvements. Overall, our results highlight the flexibility of proactive task encoding, governed by distinct representational organizations in specific brain regions. They also stress the variability of motivation-control interactions, which appear to be highly dependent on task attributes such as complexity or novelty.

**Significance Statement:** In comparison with other primates, humans display a remarkable success in novel task contexts thanks to our ability to transform instructions into effective actions. This skill is associated with proactive task-set reconfigurations in fronto-parietal cortices. It remains yet unknown, however, *how* the brain encodes in anticipation the flexible, rich repertoire of novel tasks that we can achieve. Here we explored cognitive control and motivation-related variables that might orchestrate the representational space for novel instructions. Our results showed that different dimensions become relevant for task prospective encoding depending on the brain region, and that the lateral prefrontal cortex simultaneously organized task representations following different control-related variables. Motivation exerted a general modulation upon this process, diminishing rather than increasing distances among instruction representations.

## Introduction

Humans quickly learn from instructions which elements are relevant in a context and their respective appropriate actions. These parameters are encoded proactively in our brain in an action-based code (Brass, Liefooghe, Braem, & De Houwer, 2017; Cole, Braver, & Meiran, 2017), preparing our perceptual and motor systems in advance (Cole, Laurent, & Stocco, 2013) and facilitating success in novel environments. Instructed behavior is thus critical to avoid less effective and slow trial-and-error learning, and also enables the social transmission of task procedures. There is scarce knowledge, however, about how the informational and motivational content of novel instructions organizes neural activity in a proactive manner.

Behavioral results support the role of proactive control (Braver, 2012) on instructed action (e.g. Liefooghe, Wenke, & De Houwer, 2012; see also Cole, Patrick, & Braver, 2018; Duncan et al., 2008; Luria, 1966). Recently, neuroimaging studies have revealed a link between novel instruction preparation and the fronto-parietal (FP) network (e.g. Cole, Bagic, Kass, & Schneider, 2010; Hartstra, Kühn, Verguts, & Brass, 2011; Palenciano, González-García, Arco, & Ruz, 2018). The middle (MFG) and inferior (IFG) frontal gyri, and the inferior frontal sulcus (IFS), together with the intraparietal sulcus (IPS), encode novel instruction content both in multivoxel activity patterns (Bourguignon, Braem, Hartstra, De Houwer, & Brass, 2018; González-García, Arco, Palenciano, Ramírez, & Ruz, 2017; Muhle-Karbe, Duncan, De Baene, Mitchell, & Brass, 2017) and distributed functional connectivity (Cole, Laurent, et al., 2013). Crucially, the fidelity of information encoding is linked to the intention to implement the instruction (versus mere memorization demands; Bourguignon et al., 2018; Muhle-Karbe et al., 2017) and it is also closely related to the efficiency of behavior (Cole, Ito, & Braver, 2016; González-García et al., 2017). Nonetheless, while current studies have mainly focused on decoding the upcoming target category (González-García et al., 2017; Muhle-Karbe et al., 2017), the wider organizational structure that shapes anticipatory task representation remains unknown. To study the relevant dimensions organizing novel instruction encoding, we selected three variables known to be relevant for proactive control.

Task preparation consists of a two-step process (Rubinstein et al., 2001), composed first by an abstract goal reconfiguration and second by the activation of specific stimulus-response contingencies (De Baene & Brass, 2014; Muhle-Karbe, Andres, & Brass, 2014). Our study exploited these two phases. First, in relation to the high-level task goal setting, we manipulated the integration of information within or across feature dimensions of stimuli (Rigotti et al., 2013), a variable traditionally linked to task complexity and top-down attention (e.g. Treisman & Gelade, 1980). Second, the stimulus-response reconfiguration process was manipulated by the response set complexity, requiring single or sequential motor responses. Moreover, to explore stimuli-specific preparatory mechanisms previously documented (e.g. González-García, Mas-Herrero, de Diego-Balaguer, & Ruz, 2016; Sakai & Passingham, 2003, 2006), we also manipulated the relevant target category.

Finally, cognitive control and motivation maintain an intricate relationship during task preparation (Pessoa, 2009, 2017). Reward expectation boosts cue-locked activity across the FP network (Parro, Dixon, & Christoff, 2017), and it has been recently linked to stronger anticipatory rule encoding (Etzel, Cole, Zacks, Kay, & Braver, 2016). Nonetheless, contradictory findings have also been found (Wisniewski, Forstmann, & Brass, 2018), and a comprehensive characterization of this interaction in complex, novel scenarios is still pending. Consequently, we included economic incentives in our paradigm and assessed the nature of their effect on instruction preparation. By varying these four variables (dimension integration, response-set complexity, target category, and reward), we built a set of novel, verbal instructions that were followed by healthy participants while functional magnetic imaging (fMRI) data were collected. Using Representation Similarity Analysis (RSA; Kriegeskorte, Mur, & Bandettini, 2008), we assessed the extent to which each of our control-related variables organized instruction encoding, as well as the effect of motivation upon this organization.

## Materials and methods

### Participants

Thirty-six students from the University of Granada completed the experimental paradigm inside an MRI scanner (16 women, mean age = 22.97 years, SD = 3.32 years). All of them were right-handed, with normal or corrected-to-normal vision, and native Spanish speakers. In exchange for their participation, they received between 20 and 40€, depending on their performance on the rewarded trials (see below). They all signed a consent form approved by the Ethics Committee of the University of Granada. Four participants were later excluded due to excess of head movement (> 3mm) or poor performance (<70% of correct responses).

### Apparatus, stimuli, and procedure

For the experiment, we built a set of 192 different novel verbal instructions. Each instruction referred to two independent conditions about faces or food items that could be met or not by the upcoming grids, and their associated responses (e.g.: “*If there are two women and an additional sad person, press A; if not, press L*”). The conditions in the instructions referred to several dimensions of the stimuli: gender (*woman, man*), race (*black, white*), emotion (*happy, sad*) and size (*big, small*) of faces, or kind (*fruit, vegetable*), color (*green, yellow*), form (*round, elongated*) and size (*big, small*) of food items.

Instructions were created by manipulating in an orthogonal manner (1) the ***Integration of*** *stimuli **dimensions*** (within vs. across dimensions), (2) the ***Response set*** required (single vs. sequential) and (3) the ***Category*** of the relevant stimuli that they referred to (faces vs. food). For example, the instruction “*If there is a woman and there is a man, press A; if not, press L*” involves within-dimension integration (i.e., gender), requires a single response (a left –“A”– or a right –“L”– index button press) and is face-related. On the other hand, “*If there is a fruit and a small food item, press AL; if not, press LA*” requires across-dimension integration (the type of food and its size), demands a sequence of two button presses to respond and is food-related. Instructions referred to either 2, 3 or 4 stimuli of the target grid. Equivalent trials were created for the different levels of these three variables.

In addition, we included ***Motivation*** as another variable: half of the instructions were associated with the possibility of receiving an economic reward if responses were fast and accurate while the other half were non-rewarded. To do so, we split our 192 instructions into two equivalent sets in terms of the manipulations of the other independent variables, and also regarding the specific attributes specified (e.g., the same number of instructions referring to happy faces in both groups). We counterbalanced across participants the assignment of these two halves to the rewarded and non-rewarded conditions. The reward status of each trial was indicated by a cue consisting on either a plus (+) or a cross (x) sign, in either silhouette or filled in black. We counterbalanced across participants whether they should attend to the shape (plus vs. cross) or the appearance (contour vs. filled sign) to obtain the reward information. This way, each participant had two different cues indicating each motivation condition, preventing a one-to-one mapping between reward expectation and visual cue identity, which otherwise could generate spurious confounds in further analysis.

For each instruction, we created two grids of stimuli, one that fulfilled the conditions instructed, and another one that did not. We counterbalanced them so that individual participants saw only one of the two instruction-grid pairings. All grids were unique combinations of images of 4 faces and 4 food items, which were pseudo-randomly selected from a pool of 32 pictures, composed by 16 faces pictures (8 different identities, half of them women and half men, half with happy expression and half with sad ones, half white and half black, appearing each of them in large and small sizes), extracted from the NimStim database (Tottenham et al., 2009), and 16 food pictures (8 different items, half of them vegetables and half fruits, half in green color and half in yellow, half with a round shape and half elongated, appearing each of them in large and small sizes) obtained from available sources on the internet (all of them with Creative Commons license). Upon target presentation, the responses required were always one or two sequential button presses, performed with the left (“A”) and/or right (“L”) index. The sequence of trial events is depicted in Figure 1. Each trial started with a jittered fixation point (0.5°), with a duration that ranged from 4500 to 7500ms, in steps of 500ms (mean = 5750ms). Then, a reward cue was presented (1.5 °; 2000ms), followed by the instruction (25.75°; 2500ms). Next a second jittered fixation appeared (with the same characteristics as the previous one), and the target grid (21°) was presented for 2500ms, where participants were required to respond. Afterward, a feedback symbol was presented (1.65 °; 500ms), indicating whether the participant had earned money in that trial (with a Euro symbol), whether the response was correct but no money was achieved (tick symbol) or whether the response was incorrect (cross symbol).

**Fig. 1:**
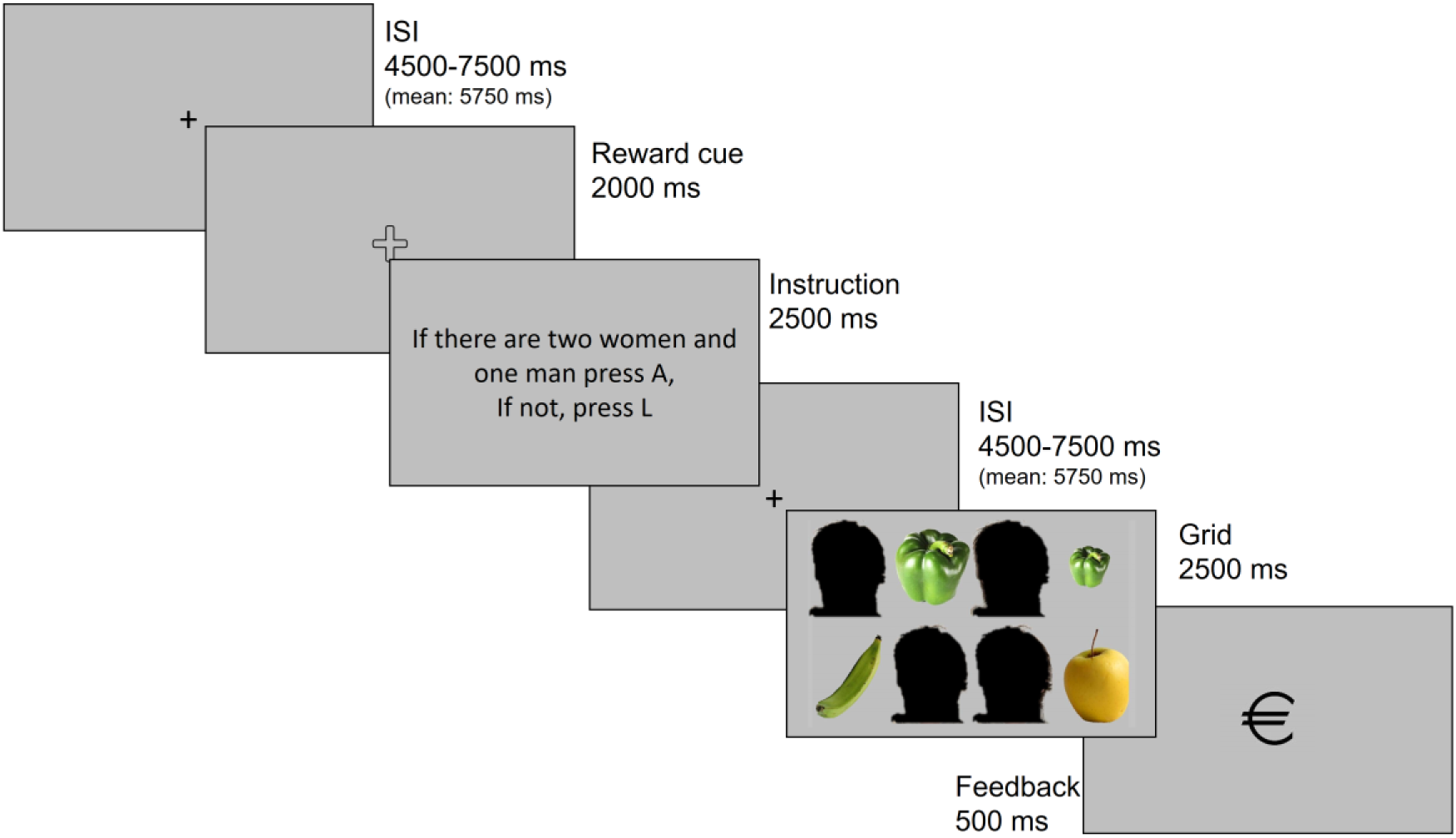
Sequence of events in a single trial. Face stimuli (obscured in the preprint version) were obtained from the NimStim database (Tottenham et al., 2009).

Before being scanned, participants completed a behavioral practice session. They received indications about how to perform the task, as well as details on how rewards would be administered, emphasizing that both accurate and fast responses were needed to accumulate money for a maximum of 40€. Specifically, they were informed that they would receive 20€ for their time and that the rest of the compensation would depend on their performance on rewarded trials: the initial extra increases would be easier to earn while approaching the upper limit of the payment would require a higher accuracy rate. Then, they performed a simple discrimination task with the different reward cues, and after that, they practiced the instruction-following task, completing one block of 32 trials. Practice instructions were drawn from a separate set (which was equivalent in all the parameters specified above) and were not employed in the MRI experiment, to maintain trial novelty. Participants repeated the practice block as many times as needed to obtain an accuracy rate above 75% (on average, participants performed the practice block 1.75 times). Once this phase was completed, the experimental paradigm was performed inside the scanner. This was composed by the full 192 instructions set, presented in six different runs (32 trials each). All runs included an equal number of face and food-related, single and sequential responses, within and across-dimension integration and rewarded and non-rewarded instructions. Overall, participants spent 90 minutes approximately inside the MRI scanner.

### fMRI preprocessing and analysis

MRI data were acquired using a 3-Tesla Siemens Trio scanner located at the Mind, Brain, and Behavior Research Center (CIMCYC, University of Granada, Spain). Functional images were collected employing a T2* Echo Planar Imaging (EPI) sequence (TR = 2210ms, TE = 23ms, flip angle = 70°). Each volume consisted of 40 slices, obtained in descending order, with 2.3mm of thickness (gap = 20%, voxel size = 3mm^3^). A total of 1716 volumes were obtained, in 6 runs of 286 volumes each. We also acquired a high-resolution anatomical T1-weighted image (192 slices of 1mm, TR = 2500ms, TE = 3.69ms, flip angle = 7°, voxel size = 1mm^3^).

The functional images were preprocessed and analyzed with SPM12 (http://www.fil.ion.ucl.ac.uk/spm/software/spm12/), with the exception of single-trial parameter estimation (see *RSA section*), which was conducted on AFNI. After discarding the first four volumes of each run to allow for stabilization of the signal, the images were spatially realigned and slice-time corrected. Then, the participants’ structural T1 image, which had been coregistered with the EPI volumes, was segmented to obtain the transformation matrices needed to normalize the functional images to the MNI space. Finally, they were smoothed with an 8mm FWHM Gaussian kernel. The full preprocessing pipeline was completed before conducting the univariate analysis, while only realigned and slice-timing corrected images were employed for the multivariate tests (see next section). In the latter, normalization and smoothing were performed after the individual-level analysis, following the same strategy as above.

### Control univariate analysis

We first conducted a univariate standard GLM, modelling each of the sixteen combinations of our variables (for example: within-dimension integration/simple response required/faces-related/ rewarded) and specifying two regressors per trial: one for the encoding phase (from the reward cue until the end of the instruction), and another for the implementation stage (encompassing the target grid presentation and until the end of the feedback cue). All regressors were convolved with the canonical hemodynamic response function. We also added error trials and six motion parameters as nuisance regressors, and a high-pass filter of 128s to avoid low-frequency noise.

The rationale of this analysis was to check the effect of motivation during the encoding of novel instructions with the aim of ensuring that our manipulation successfully generated typical reward-related patterns of activation (Parro et al., 2017). This was done by performing *t-*tests at the individual (first) level, contrasting rewarded versus non-rewarded encoding regressors, and carrying these statistical maps to a group one-sample *t-*test. The result was cluster-wise FWE-corrected for multiple comparison at *P* < .05 (from an initial threshold of *P* < .001 and k = 10). With this approach, we obtained one large cluster that extended across multiple brain regions. To obtain smaller, anatomically coherent clusters, we employed a stricter threshold (uncorrected cluster-forming threshold of *P* < .0001, with the corresponding FWE correction at *P* < .05), as done previously (e.g. Dumontheil et al., 2011; Palenciano et al., 2018).

### Representational Similarity Analyses

We conducted a series of multivariate RSAs, following a two-step approach. First, we analyzed whole-brain data, using a searchlight approach, to find regions encoding novel instructions according to each of our three control-related variables. Second, we used the significant areas as Regions Of Interest (ROIs) and focused on them to explore the effect of reward on their representational geometry.

### Whole-brain model-based RSA

We first studied whether the representational structure of novel instructions was explained by three variables related to cognitive control preparation: dimension integration, response set complexity and target category. Importantly, we specifically wanted to explore this during the initial encoding stage, where proactive task-set reconfiguration takes place. To do so, we first obtained trial-by-trial estimations of our signal, following a Least-Square-Sum approach (LSS; Turner, 2010) to ensure the smallest possible collinearity among regressors (Arco, González-García, Díaz-Gutiérrez, Ramírez, & Ruz, 2018). We generated and estimated one separate model per trial, in which we defined: (1) a regressor isolating the encoding phase of the individual trial of interest; (2) a second regressor containing the rest of trials (encoding phase) of the same condition; (3) thirty-one additional regressors encompassing the rest of conditions at the encoding and implementation phases (as in the GLM specified above), and (4) nuisance regressors (movement, errors). To do so, we employed AFNI’s function 3dLSS (https://afni.nimh.nih.gov/pub/dist/doc/program_help/3dLSS.html). Once the trial-wise parameter images were obtained, the rest of the RSA was performed with The Decoding Toolbox (Hebart, Görgen, & Haynes, 2014).

In our analysis, we compared three theoretical models of representational organization (one per preparation-related independent variable) with the empirical one, built from spatially distributed activity patterns. To do so, we employed a spherical searchlight (radius: 4 voxels) and applied it to the whole brain (Kriegeskorte, Goebel, & Bandettini, 2006). First, we built three theoretical representational dissimilarity matrices (RDM, Fig. 2a), which captured the expected dissimilarity (represented with 0s and 1s) between pairs of trials, according to the corresponding variables of interest. For example, in the Category RDM, dissimilarity is expected to be minimal within pairs of trials that refer either to faces or to food, while maximal between pairs of trials referring to different target categories. Then, in each iteration of the searchlight, we generated a neural RDM, using a measure of distance based on Pearson correlation. Specifically, we extracted the corresponding single-trial beta values of the voxels involved, correlated each pair of the trials’ activity patterns, and subtracted that value from 1. Afterwards, this neural RDM was Spearman-correlated with the theoretical ones (Fig. 2c), and the coefficients were normalized with Fisher’s z transformation and assigned to the central voxel of the searchlight sphere. Importantly, both theoretical and neural matrices were built trial-wise (i.e., not averaging within conditions), and thus, were fully symmetrical with a diagonal of 0s. Consequently, only the lower triangle of the matrices, excluding the diagonal, was included in the correlation to avoid inflated positive results (Ritchie, Bracci, & Op de Beeck, 2017). After iterating the searchlight across the whole brain, we obtained three maps per participant representing how well the representational geometry in different regions matched the one expected by each of our three theoretical models.

**Fig. 2:**
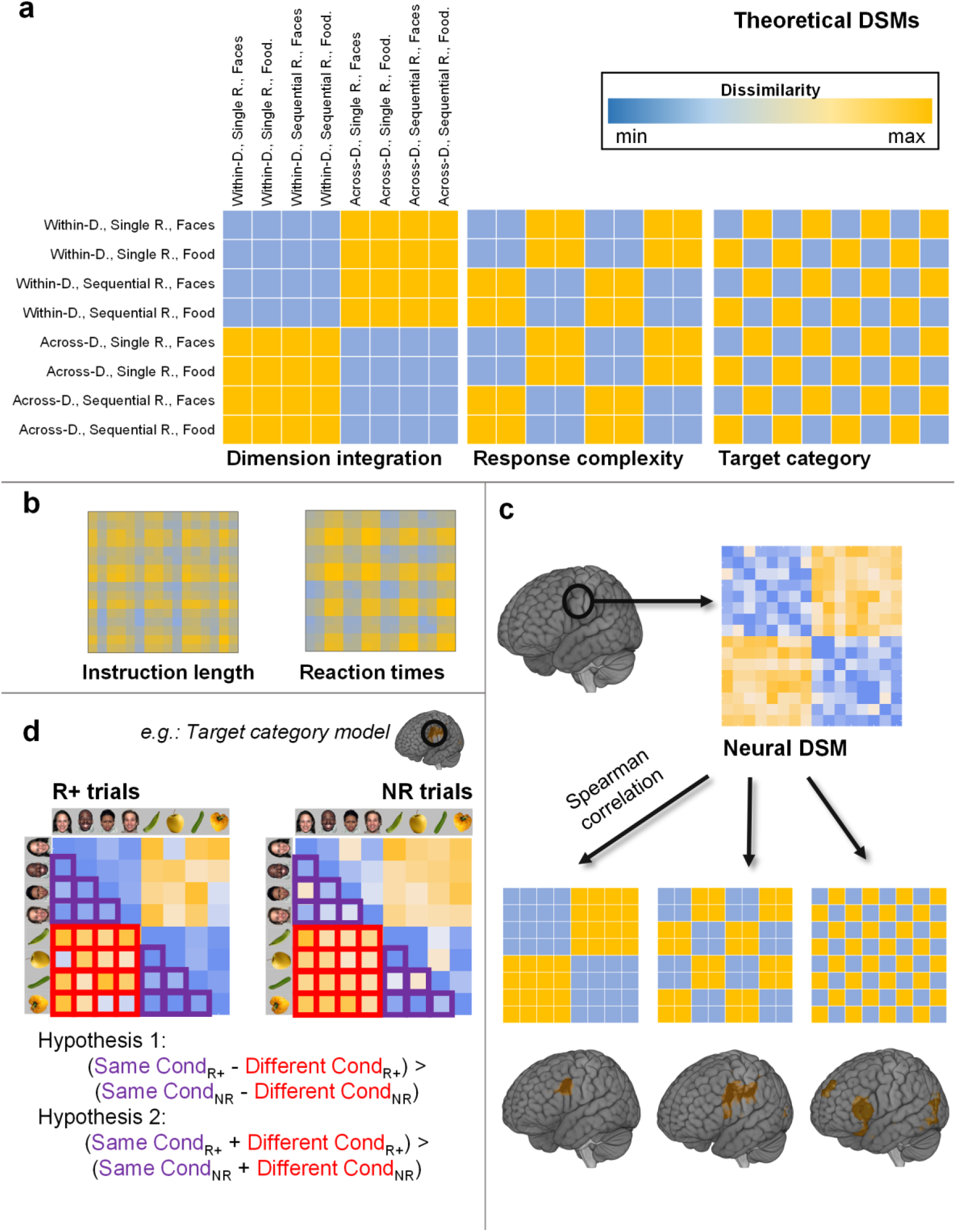
Main analysis procedure. (a) Theoretical Representational Dissimilarity Matrices (RDMs) employed in the Representational Similarity Analysis (RSA). Within/Across-D. stands for within-dimension and across-dimension integration, while Single/Sequential R. stands for single response and sequential response. (b) RDMs capturing differences in instruction length (number of letters) and reaction time, included in a multiple regression analysis together with matrices shown in (a) to control for the effect of these two variables. (c) Following a searchlight approach, we extracted the neural RDM at each brain location and compared it – via Spearman correlation – with our three theoretical RDMs. As a result, we obtained three whole-brain correlation maps, one per model. (d) To assess the effect of motivation, for each region significant in (c) we extracted the neural RDMs from rewarded (R+) and non-rewarded (NR) trials. To study potential interactions of reward expectation and the corresponding model variable (Hypothesis 1), we averaged the dissimilarity values among same-condition and different-condition trials and tested if the subtraction among these two values was higher in the rewarded condition (using Wilcoxon signed-rank test). We also checked for a general increase in dissimilarities associated to reward (Hypothesis 2). *Note*: All matrices in the figure were simplified for visualization purposes by averaging cells within conditions. The matrices shown in (b) were further averaged across the sample. In (d), matrices display only one task variable (collapsing between the remaining two) to highlight the analysis logic. In all the analyses, however, trial-wise and single subject matrices were employed.

Statistical significance was assessed non-parametrically via permutation testing, as proposed by Stelzer, Chen, & Turner (Stelzer, Chen, & Turner, 2013). We first performed 100 permutations at the individual level, where trial labels were randomly shifted and the whole analysis was repeated. Then, at the group level, we resampled 50,000 times one of the permuted maps of each subject and averaged them. The resulting bootstrapped group maps were used to build a voxel-wise null distribution of correlation values, which was used to extract the correlation coefficient coinciding with a probability of 0.001 of the right-tailed area of the distribution (i.e., linked to a p <= .001) of each individual voxel. The group map of the results was then thresholded using these values. From the bootstrapped maps we also built a null distribution of cluster sizes (Stelzer, Chen, & Turner, 2013), which determined the probability of each cluster extent under the null distribution. We used this to assign the corresponding *P* value to the surviving clusters of the group results map, and FWE-corrected (P < .05) them to control for multiple comparisons.

We performed a further conjunction test to find areas sharing the three representational organizational schemes. To do so, we thresholded (*P* < .05, FWE corrected) and binarized the three maps from the previous step, and obtained the overlapping voxels (Nichols, Brett, Andersson, Wager, & Poline, 2005).

Importantly, the RSA results could be influenced by other variables statistically related to our manipulations (Popov, Ostarek, & Tenison, 2018), such as instructions’ length and speed of responses, which differed slightly between conditions. To examine their influence on the results, we performed an additional multiple regression analysis taking both variables into account. We built two different RDMs (see Fig. 2.b) in which each cell contained the absolute difference in the number of letters (instruction’s length RDM) or reaction time (response speed RDM), respectively, between specific pairs of instructions. We then used them as regressors together with the three proactive control-related RDMs, predicting the neural pattern of dissimilarities in each iteration of a searchlight. The regressors were built vectorizing the lower triangle of the RDM, excluding the diagonal values. It is important to note that there were small but still significant correlations among some of the regressors included in the analysis. Specifically, dimension integration correlated with instruction length and RT, and target category did so with instruction length. To assess the impact of these correlations on the regression estimation, we computed Variance Inflation Factors (Mumford, Poline, & Poldrack, 2015), an index of the regressors’ collinearity. For our five models, and in all the participants, VIF were always below 1.1 (being 5 a typical cutoff above which the estimation would be compromised; Mumford et al., 2015). Thus, even despite the relationship among variables, the results of our main analyses are still meaningful. The corresponding beta weight maps obtained showed the regions where the effect of our variables of interest remained significant even when instruction’s length and response speed were included.

Finally, even when the distance measure employed to build the neural RDMs (i.e., Pearson correlation) is insensitive to differences in mean signal intensity between conditions, differences in signal variance could be affecting it (Walther et al., 2016). For that reason, these analyses as well as the reward-related tests (see below), were repeated after a z-normalization of the multivoxel activity patterns, ensuring equal mean (0) and standard deviation (1) across all pairs of trials. The results thus obtained did not differ from the initial non-normalized ones, so we do not report them here.

### ROI-based RSA

The previous analysis identified brain areas encoding instructions according to each one of three proactive control variables, separately. We next ran ROI analyses to further explore the role of the three variables for task coding in these regions. Specifically, we estimated the extent to which each of the manipulated control variables explained the neural organization in the ROIs identified in the previous analysis. We followed a Leave-One-Subject-Out (LOSO) cross-validation procedure (Esterman, Tamber-Rosenau, Chiu, & Yantis, 2010), using the searchlight maps obtained before. First, we identified regions sensitive to each of the three models for each participant, running a group level *t*-test with the corresponding maps from the rest of the sample, i.e., excluding their own data. Significant clusters showing consistency across all LOSO iterations were selected as ROIs, and inverse normalized to the participants’ native space. In a second step, we estimated the ROIs RDMs and correlated them with the three models RDMs. Importantly, thanks to the LOSO procedure we avoided circularity in the analysis, as independent data was employed to select the ROIs and to compute de correlations with the models. The correlation coefficients (for each participant, one per ROI and model) were then introduced in a repeated measures ANOVA, with ROI and Model as factors, and the interaction term was examined to detect heterogeneity in task encoding organization across regions (Reverberi, Gorgen, & Haynes, 2012). Interactions were further characterized by one sample *t*-tests, in order to determine which models had an effect on the different regions studied. Whenever the normality assumption was not met (assessed with the Saphiro-Wilk test), we employed Wilcoxon signed-rank tests instead. All *P* values were Bonferroni-corrected for multiple comparisons, adjusting them to the number of ROIs explored.

Additionally, we aimed to extrapolate our findings to regions consistently found in the literature during both practiced (e.g. Woolgar, Hampshire, Thompson, & Duncan, 2011) and novel (e.g. González-García et al., 2017) task preparation, and in general, when demanding cognitive processing is deployed (Duncan, 2010). This set of brain areas belong to the Multiple Demand Network (MDN; Duncan, 2010), which includes the bilateral RLPFC, MFG, IFS, anterior insula/frontal operculum (aIfO) area, IPS, anterior cingulate cortex (ACC) and pre-supplementary area (preSMA). To assess the organization of novel task encoding across this MDN, we employed functionally derived masks of its nodes (from Fedorenko, Duncan, & Kanwisher, 2013; template available at http://imaging.mrc-cbu.cam.ac.uk/imaging/MDsystem), inverse normalized them to the participants’ native space, and followed the same ROI-approach as above, extracting each ROI RDM and correlating it with the models’ matrices. Again, correlation coefficients were entered into a repeated measures ANOVA with ROI and Model as factors, interactions were examined, and finally, a series of one-sample *t*-tests (or Wilcoxon signed-rank test when normality was violated) were conducted.

### Analysis of reward-related effects on RSA results

A final goal of our study was to assess whether the representational space of novel instructions was affected by motivation. Our initial hypothesis was that reward would polarize the representational geometry, enhancing the effect of our control-related variables at structuring rule encoding. In other words, and taking as an example the target category variable, we assessed whether reward expectations would increase the distance between representations of instructions referring to different stimulus categories (in extension to the other variables, indicated as *different-condition dissimilarity*), while decreasing the distance among those referring to same target category (*same-condition dissimilarity*). Our second, alternative hypothesis was that reward would exert a general effect, globally increasing the distances among instruction representations, independently of the other variables manipulated. In this sense, we expected that both *different* and *same-condition dissimilarity* would be increased in rewarded trials, in comparison with non-rewarded ones. The two possibilities would be compatible with previous findings showing that reward expectancy enhances rule decodability (Etzel et al., 2016).

To test these two hypotheses, we run ROI analyses (Fig. 2d) for each of our control-related variables, focusing on the regions that resulted statistically significant in the main RSA. To do so, at the individual level and for each variable, we first ran a searchlight and generated four whole-brain maps containing dissimilarity values among: (1) same-condition rewarded trials; (2) different-conditions rewarded trials; (3) same-condition non-rewarded trials; and (4) different-conditions non-rewarded trials. These values were the result of averaging and normalizing (with the Fisher transformation) the pertinent cells of the neural RDM (see Fig. 2c for an example) in each searchlight iteration. The maps thus obtained were normalized to the MNI space, so we could extract participants’ mean dissimilarities for each of our ROIs using MarsBar (Brett, Anton, Valabregue, & Poline, 2002). After that, and for each ROI and variable, we conducted two Wilcoxon signed-rank tests (Nili et al., 2014). First, to assess our main hypothesis, we tested whether (DifferentCond._Rewarded_ - SameCond._Rewarded_) > (DifferentCond._NonRewarded_ - SameCond._NonRewarded_). To explore the second possible hypothesis, we collapsed across same and different conditions, and tested if (DifferentCond._Rewarded_ + SameCond._Rewarded_)/2 - (DifferentCond._NonRewarded_ + SameCond._NonRewarded_)/2 was greater than 0 (Fig 2c). In both analyses, we corrected for multiple comparisons (number of ROIs being tested) with an FWE threshold of *P* < .05.

Last, to investigate the relevance for behavior of the effect of motivation on representational structure, we correlated this effect with behavioral data. Specifically, for each participant, we computed the average decrease in dissimilarity and in the inverse efficiency scores (IES; Townsend & Ashby, 1978) linked to rewarded trials (in comparison with non-rewarded ones). The IES was employed in this analysis to take into account, simultaneously, improvements in accuracy and response speed. As we performed as many correlations as ROIs assessed in this analysis, we again controlled for multiple comparisons with an FWE threshold of *P* < .05. Additionally, to explore the possibility of motivation exerting an effect during the subsequent implementation of instructions, we also ran the analyses detailed above with beta images obtained from this stage.

### MVPA-based assessment of reward effects

Finally, to further connect our results with previous findings, we performed multivoxel pattern analysis (MVPA) to explore the effect of reward on decoding precisions (Etzel et al., 2016). We decoded the two conditions of each of our three control-related variables, training three binary classifiers: one for distinguishing between within versus across-dimension integration instructions, other for single versus sequential response requirements, and the last one for faces and food-related trials. This was done separately for rewarded and non-rewarded trials. Again, we used non-normalized and unsmoothed trial-wise beta images from the encoding stage. As we aimed to detect any region with reward-related increases in task decodability, we performed the MVPA in a whole brain fashion, using searchlight (instead of biasing the results using ROIs resulting from the RSA). In each searchlight iteration, we followed a leave one-run-out cross-validation approach, training a linear support-vector machine classifier (C=1; Pereira, Mitchell, & Botvinick, 2009) with five of our six runs, and testing it with the remaining one, in an iterative fashion. Then, for each of our variables, we subtracted the accuracy map obtained from non-rewarded trials to the map from rewarded ones, and then normalized and smoothed these images, to conduct an above zero one-sample *t*-test at the group level. This way, we assessed the benefits in classification precision associated with reward.

## Results

### Behavioral results

We analyzed RT and accuracy data separately, conducting two repeated measures ANOVA with four factors, corresponding to the four variables manipulated: dimension integration (within vs. across), response set complexity (single vs. sequential), category (faces vs. food items) and motivation (rewarded vs. non-rewarded). Importantly, the main effect of motivation was statistically significant on both accuracy (F_1, 31_ = 4.97, *P* < .05, η_*p*^2^_ = .14) and RT (F_1, 31_ = 6.52, *P* < .05, η_*p*_^2^ = .17) data, with more accurate (rewarded: M = 0.85, SD = 0.11; non-rewarded: M = 0.83, SD = 0.12) and faster (rewarded: M = 1.16, SD = 0.21; non-rewarded: M = 1.20, SD = 0.20) responses on the rewarded condition (see Fig. 3). This indicates that participants made use of reward cues and the economic incentives had the expected effect on behavior, improving its efficiency

**Fig. 3:**
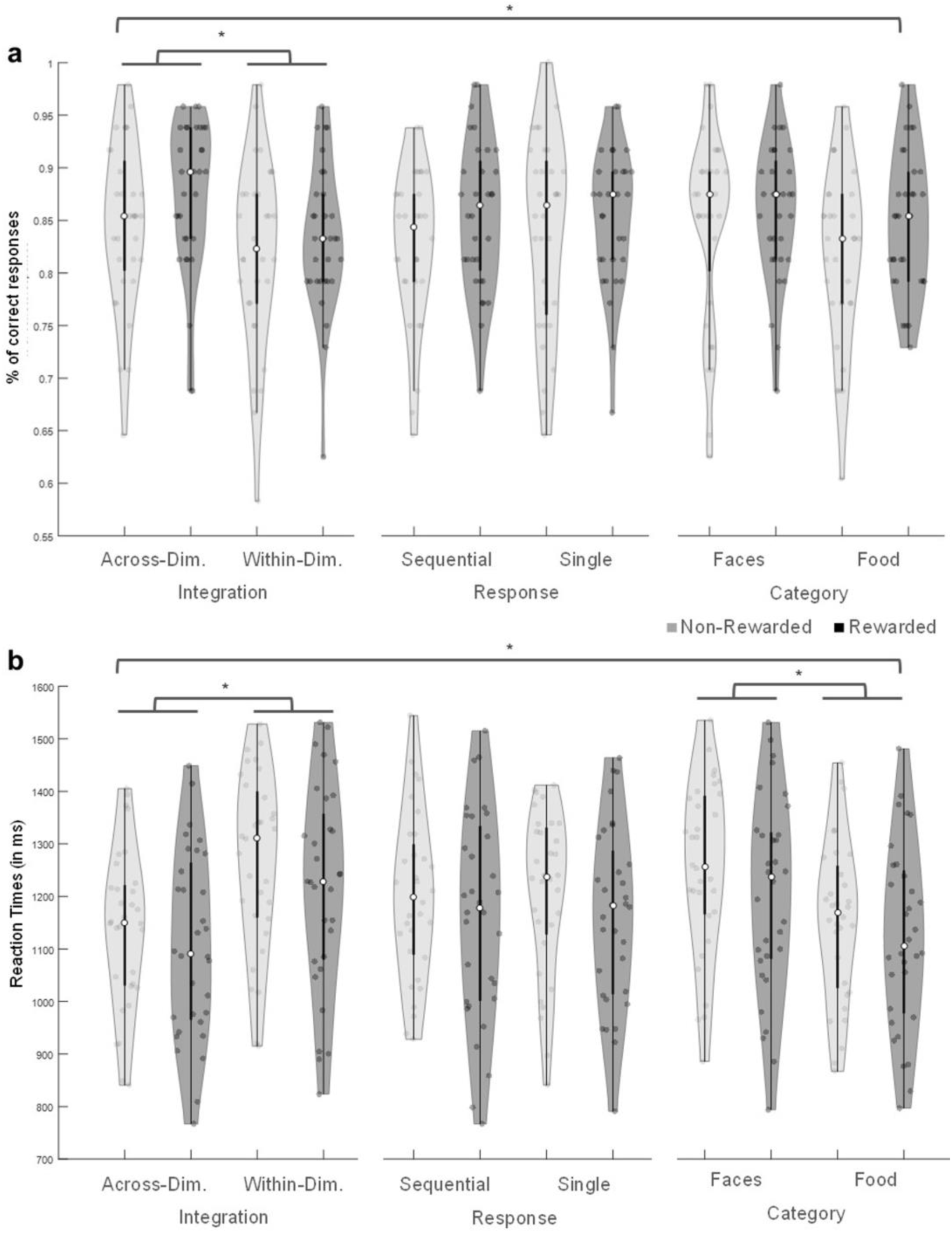
Behavioral data. Violin plots showing correct responses (a) and Reaction Time (b) data for each condition, in rewarded and non-rewarded trials.

In addition, accuracy data showed a main effect of dimension integration (F_1, 31_ = 9.24, *P* < .05, *η_p_*^2^ = .23), with better performance when within-dimension integration was required (within dimension: M = .86, SD = 0.13; across dimensions: M = .83, SD = 0.12), and a significant three-way interaction of category, response set complexity and dimension integration (F_1, 31_ = 4.46, *P* = .043, *η_p_*^2^ = .13). Even despite the lack of hypothesis regarding an interaction at this level, we performed post hoc pair-wise comparisons, which revealed that the interaction was driven by less robust (*P* > .05) differences among within and across-dimensions trials that required a single response and was food-related (while, in the rest of combinations of independent variables, this difference was significant).

On the other hand, RT results also showed a main effect of dimension integration (F_1, 31_ = 61.81, *P* < .001, *η_p_*^2^ = .67) in the same direction as above (within-dimension: M = 1.12, SD = 0.17; across-dimensions: M = 1.24, SD = 0.2), and a main effect of category (*F*_1, 31_ = 74.89, *P* < .001, *η_p_*^2^ = .71), with faster responses to food-related instructions (faces: M = 1.23, SD = 0.21; food items: M = 1.14, SD = 0.19). Neither the effect of response set complexity (accuracy: F_1, 31_ = 0.31, *P* = .579, *η_p_*^2^ = .01; reaction time: F_1, 31_ = 0.21, *P* = .653, *η_p_*^2^ = .01) nor any other ANOVA term resulted significant in the behavioral measures (main effect of Category on accuracy: F_1, 31_ = 3.23, *P* = .082, *η_p_*^2^ = .094; all interactions terms, except the ones stated above, P > .100).

### Univariate results: reward-related activations during instruction encoding

We first assessed mean activity during novel instruction encoding, comparing rewarded against non-rewarded trials. To do so, we performed a univariate GLM, defining regressors for each combination of variables (e.g.: within-dimension integration, single response, face-related rewarded trials), separately for the encoding and the implementation stages. A group level *t*-test showed that, in accordance with our expectations and previous literature (Parro et al., 2017), the basal ganglia and fronto-parietal cortices were more active for rewarded than non-rewarded instruction encoding. We observed peaks of activation (see Fig. 4) in the bilateral inferior frontal junction (IFJ), premotor and supplementary motor areas (left: [-33, 5, 26], z = 5.07, k = 489; right: [33, 2, 59], z = 4.79, k = 572), cingulate cortex ([-9, 5, 32], z = 5.48, k = 20), bilateral IPS extending into the precuneus (left:[−18, −64, 35], z = 4.77, k = 357; right: [33, −52, 53], z = 4.36, k = 324), the accumbens, ventral portion of the caudate and thalamus ([12, −22, 20], z = 5.13, k = 1176), inferior temporal gyrus ([48, −58, −13], z = 4.48, k = 52), occipital cortex ([30, −61, −25], z = 5, k = 1364) and midbrain ([0, −31, −4], z = 5.19, k = 255). Thus, regions involved in reward processing (Haber & Knutson, 2009), as well as in cognitive control paradigms with monetary incentive manipulations (e.g. Engelmann, 2009), were engaged by our task, indicating the success of the reward manipulation.

**Fig. 4:**
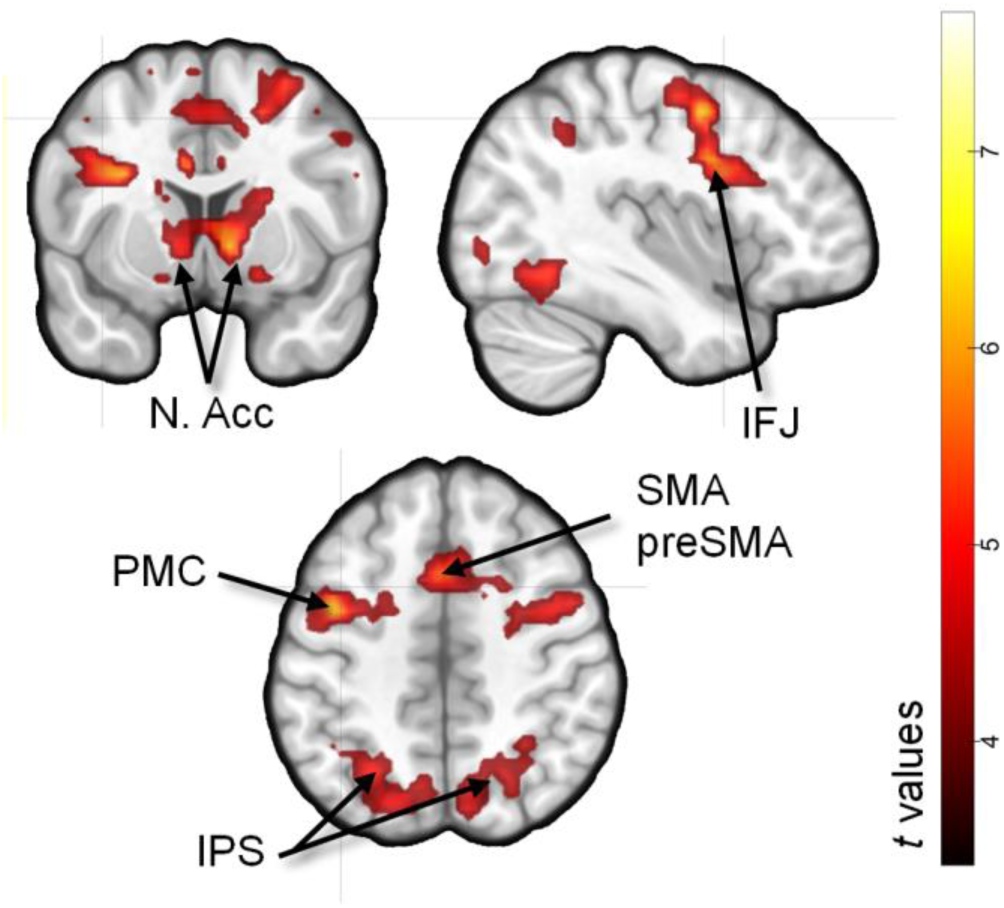
Regions showing greater activity during the encoding of rewarded compared to non-rewarded instructions. Abbreviations stand for Nucleus Accumbens (N. Acc), inferior frontal junction (IFJ), premotor cortex (PMC), supplementary motor cortex (SMA), pre-supplementary motor cortex (preSMA) and intraparietal sulcus (IPS).

### Model-based RSA results: instruction encoding structured by proactive-control variables

We aimed to identify regions whose organization during task encoding was explained by dimension integration, response set complexity and target category. With that purpose, we employed an RSA (Kriegeskorte, Mur, & Bandettini, 2008) to compare the representational dissimilarity matrices (RDMs) found in neural data during the encoding stage with theoretical RDMs corresponding to the three proactive control-related variables (see Fig. 2). In neural RDMs, each cell contained the dissimilarity (1 – Pearson correlation) between the multivariate patterns of activation of two trials. In the theoretical RDMs, cells contained dissimilarities (1: maximal, 0: minimal) that we would expect if a certain variable organized encoding (i.e.: for target category, all faces-related trials would be minimally dissimilar, while face and food-related trials would be maximally dissimilar). Using searchlight (Kriegeskorte et al., 2006), we Spearment-correlated neural and theoretical RDMs across the brain and obtained maps showing how well these three variables captured the representational space of different areas. The modality of **dimension integration** (Fig. 5a) only had a significant effect on rule encoding at the left MFG and IFG, incurring into the IFS ([-51, 20, 26], k = 642). **Response set complexity** (Fig. 5b), on the other hand, organized task representations on a wide cluster including the bilateral IFG, premotor, supplementary and primary motor cortices, somatosensory area, middle temporal gyrus and superior and inferior parietal lobe extending along the IPS ([-42, −31, 44], k = 8583) and in the left parahippocampal cortex ([-18, −40, −1], k = 301). Finally, in the case of the **target category** RSA (Fig. 5c), significant correlations were found in an extensive cluster on the left hemisphere covering the IFG incurring into the IFJ, the fusiform gyrus, the temporo-parietal junction (TPJ), the inferior and middle temporal gyrus and the precuneus ([-39, −67, 17], k = 5581). On the right hemisphere, the analysis was also significant on the right middle temporal gyrus and TPJ ([39, −58, 23], k = 442) and the IFG ([42, 26, 14, k = 295]. Finally, the medial superior frontal gyrus ([-9, 53, 26], k = 377) was also involved.

**Fig. 5:**
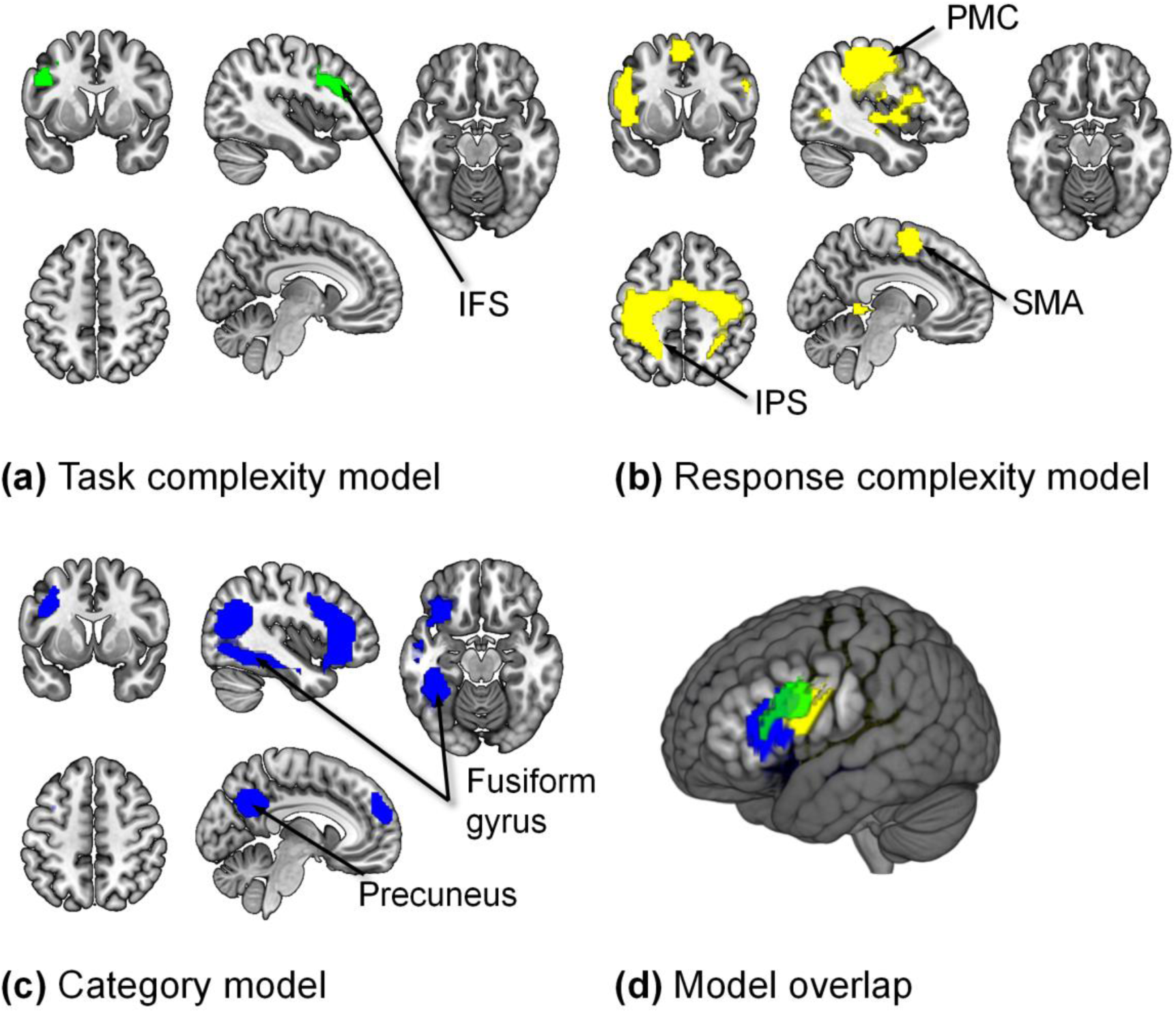
Model-based RSA searchlight results for the three models (a-c) and render image showing the overlap among them (d). *Note:* Identical sections were employed to display the results across models.

As instructions’ length and speed of responses varied among some of our variables, we performed an additional multiple regression analysis, in which we included our three theoretical models, an RDM based on dissimilarities in length, and another one based on RT as regressors. Importantly, the multiple regression statistical model was examined to detect an excess of collinearity which could have impaired the interpretability of these results. We computed the VIF for all the regressors and across our whole sample of participants, and all of were under 1.1, an index of good estimability of regression weights. The beta maps (one per model) obtained after iterating the analysis in a searchlight procedure ensured that the variance linked to our RSA models was not misattributed due to differences in instruction length or speed of responses. Importantly, the results obtained this way were very similar to the ones extracted with the standard approach, identifying the same clusters than before. We also conducted a **conjunction analysis** to assess the overlap among regions common to the three organizational schemes. Only the left IFG and IFJ resulted significant in this test (Fig. 6).

### LOSO-based ROI analysis: assessing confluence of models within regions

The previous analyses left unexplained the extent to which each of the brain areas isolated by RDM analyses reflected in their organization the three manipulated variables. Furthermore, the conservative correction for multiple comparisons used in the searchlight could overshadow this effect elsewhere in the brain. To shed some light upon this issue, we employed a more sensitive ROI analysis, together with a LOSO approach to avoid double dipping when selecting regions.

**Fig. 6:**
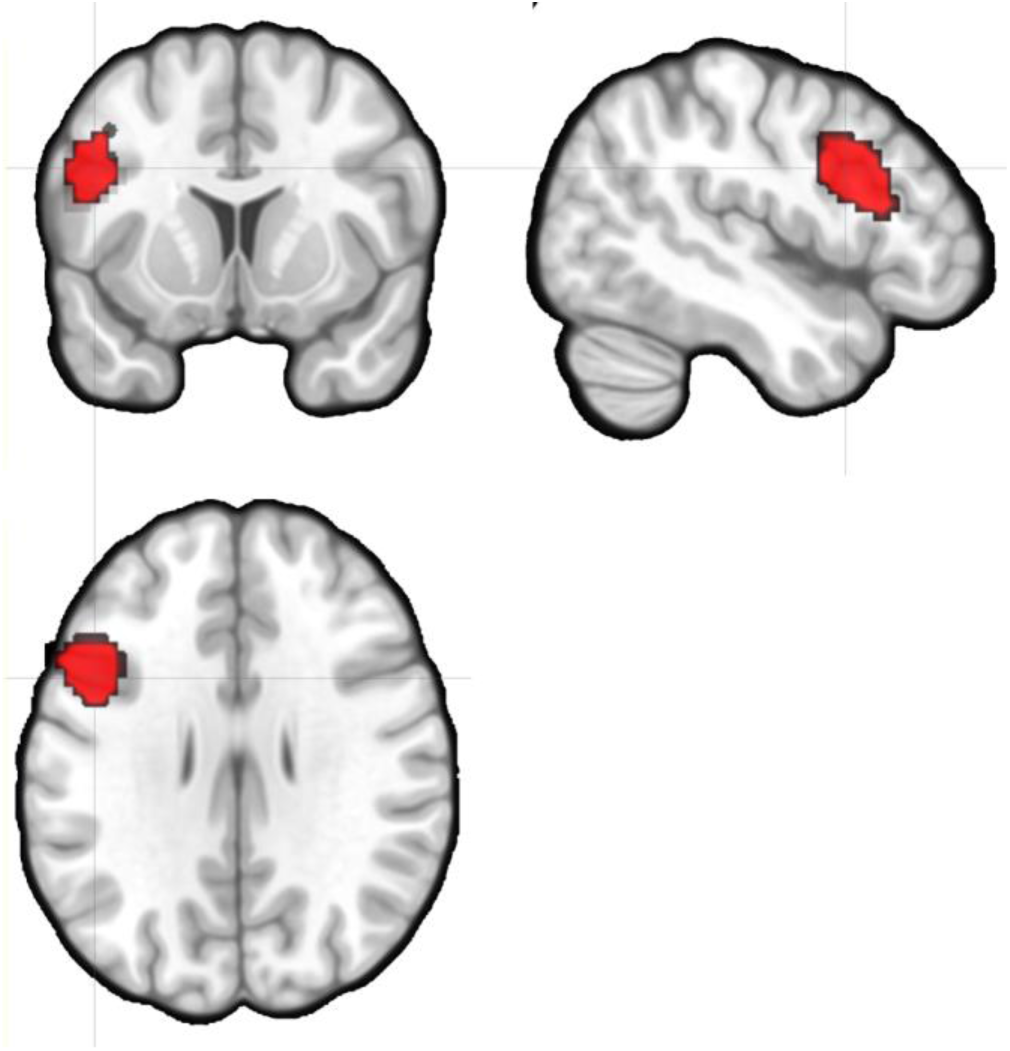
Conjunction analysis results.

All the clusters identified in the main group results (Fig. 5) were consistently found across all participants with the LOSO approach, with the exception of the medial superior frontal gyrus under the category model, which was absent in four subjects and thus not included in the analysis. The correlations of the ROIs’ RDMs and the three models’ matrices were analyzed with a repeated measures ANOVA, in which we found a significant interaction of ROI and Model (F12, 348 = 6.050, *P* < .001, *η_p_^2^* = .173), evidencing variability in instruction coding structure across regions. We then ran one sample *t*-tests or Wilcoxon signed-rank tests (depending on data distribution) to assess model performance in each ROI (see Table 1). The general pattern obtained replicated the searchlight results: the model which originally identified each specific ROI in the searchlight was the one explaining most robustly its encoding activity. Further, in almost all the regions, we did not find enough evidence supporting the effect of the remaining variables. Converging with the previous analyses, the left IFG identified with the dimension integration model was also significantly correlated with response set complexity and category. Similarly, the left IFG cluster found in the category RSA was correlated with the dimension integration model too. In addition, this confluence of models analysis revealed that the response set model was also significant in the category-related cluster involving the left fusiform and precuneus (see Table 1).

**Table 1.** Effect of the three models on the LOSO-estimated ROIs.

| Original model | ROI | Model tested | T value | Z value | P value |
| --- | --- | --- | --- | --- | --- |
| Dimension integration | Left IFG | Dim. | 3.354 |  | .008 |
|  |  | Resp. | 3.292 |  | .009 |
|  |  | Cat. | 3.635 |  | .004 |
| Response set complexity | Left IPS | Dim. | 0.614 |  | 1 |
|  |  | Resp. | 5.351 |  | < .001 |
|  |  | Cat. |  | 1.975 | .163 |
|  | Motor cortices, left LPFC | Dim. | 2.478 |  | .067 |
|  |  | Resp. | 3.647 |  | .004 |
|  |  | Cat. | 1.166 |  | .886 |
| Target category | Left fusiform gyrus and precuneus | Dim. | 0.476 |  | 1 |
|  |  | Resp. | 3.463 |  | .006 |
|  |  | Cat. | 5.466 |  | < .001 |
|  | Left IFG | Dim. | 2.832 |  | .029 |
|  |  | Resp. |  | 0.699 | .242 |
|  |  | Cat. | 4.930 |  | < .001 |
|  | Right MTG | Dim. |  | -0.144 | .557 |
|  |  | Resp. |  | -1.008 | .843 |
|  |  | Cat. |  | 2.859 | .002 |
|  | Right IFG | Dim. |  | 1.275 | .101 |
|  |  | Resp. |  | -0.206 | .582 |
|  |  | Cat. |  | 3.085 | .001 |

Note: P values displayed are Bonferroni-corrected for multiple comparisons. Abbreviations stand for inferior frontal gyrus (IFG), intraparietal sulcus (IPS), and middle temporal gyrus (MTG), Dimension integration model (Dim.), Response complexity model (Resp.) and Target Category (Cat.).

### ROI analysis spanning Multiple Demand Network regions

Following a similar strategy as above, we also examined task encoding organization across the regions comprising the MD network. We extracted each MD region’s RDM and correlated it with our three models’ RDM, and then entered the correlation coefficients into a repeated measures ANOVA. Again, a significant ROI*Model interaction was found (F_20, 620_ = 2.168, *P* = .002, *η_p_^2^* = .065). To assess which models significantly structured activations across MD ROIs, we conducted one-sample *t*-tests or Wilcoxon signed-rank tests when data were not normally distributed (see Table 2).

**Table 2.**
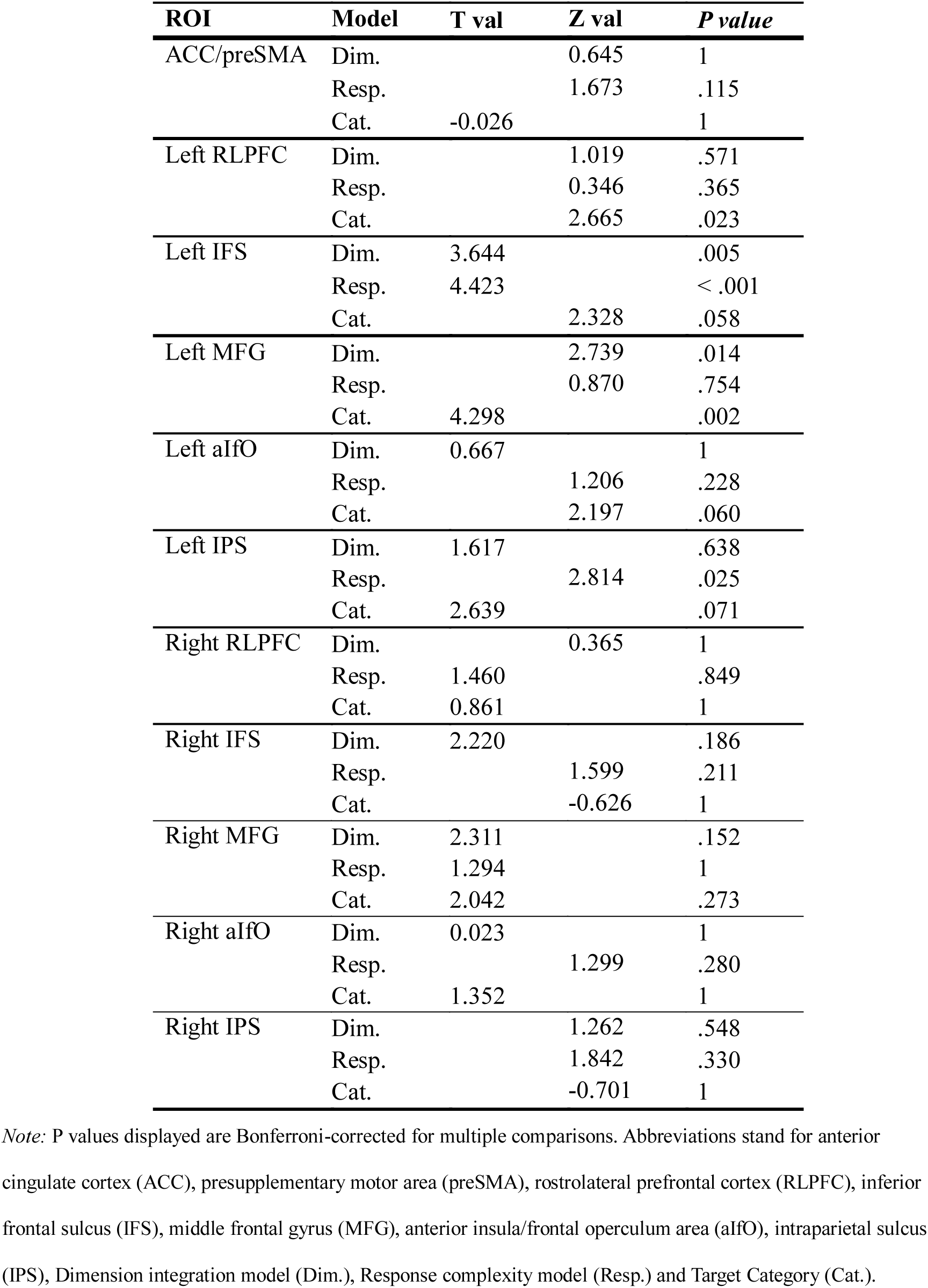
Effect of the three models on the MD network ROIs.

Only a subset of MD network regions encoded instructions consistently according to any of the proactive control variables, and all of them were located on the left hemisphere and in the LPFC and parietal cortex. The findings were, however, consistent with the searchlight and ROI-related results presented so far. The three variables exerted an effect on different left lateral prefrontal sections: dimension integration and response complexity on the IFG; dimension integration and target category on the more dorsal MFG; and finally, category on the RLPFC. Response complexity was the attribute which most robustly captured representational organization in the IPS.

### Effects of reward on representational geometry

We then explored the effects of motivation in each of the ROIs encoding different attributes of the instructions (Fig. 5), assessing two possible mechanisms that could underlie the behavioral improvements linked to reward (Fig. 2). On the one hand, we tested whether reward made our variables more efficient in sharpening the representational space (Fig. 2d, Hypothesis 1), In other words, and taking as an example the target category variable, we assessed whether reward expectations would increase the distance between representations of instructions referring to different stimulus categories (in extension to the other variables, indicated as *different-condition dissimilarity*), while decreasing the distance among those referring to same target category (*same-condition dissimilarity*). On the other, we tested the alternative possibility that dissimilarities would be, in general, greater in the rewarded trials (Fig 2d, Hypothesis 2), regardless of the variables manipulated (i.e., regardless of the pair of instructions being same or different-condition). This could reflect a mechanism for making rule representations more distinguishable among each other, and also, it would be compatible with the increase in rule decoding accuracy that has been liked to motivation in previous reports (Etzel et al., 2016). With that purpose, we extracted, for each region, the average dissimilarity among pairs of instructions pertaining to the same and different conditions, separately for rewarded and non-rewarded trials. We then used Wilcoxon signed-rank tests (Nili et al., 2014) to check whether the difference between different-condition and same-condition trials was larger in the rewarded than in the non-rewarded condition, and also, whether the mean dissimilarity (collapsing across same and different-condition) was increased by motivation.

In the first case, no reward-related differences were observed for any of the instruction-related variables (all *P*s >.1). It is important to note, however, that these results (as most of the findings presented in this study) are anchored to the instruction’s encoding stage, in which proactive control configuration takes place. To explore the possibility that the hypothesized interaction shaped neural activations during the later implementation phase (more related to reactive control; Braver, 2012; Palenciano, González-García, Arco, & Ruz, 2018), we conducted a further test employing beta images from this epoch. However, and again, the expected effect was not significant for any of the ROIs examined (all *P*s >.1).

When addressing the second hypothesis, surprisingly, we found the opposite pattern: reward systematically decreased the dissimilarity values in all the ROIs evaluated (all *P*s < .05, see Table 2). To test the behavioral relevance of this finding we correlated, across our participants, the average decrease in dissimilarities associated with reward, with the benefit of motivation on performance (IES; Townsend & Ashby, 1978). We found that in fact, the decrease in representational distances due to reward was significantly correlated with the motivation-related improvements in behavioral performance. Furthermore, this seemed to be a quite robust effect, being present in all of the ROIs included in the analysis (see Table 3 for further details).

**Table 3.** Effect of reward on dissimilarity values and correlation with behavioral improvement.

| ROI | Effect of reward on<br>dissimilarity values | Correlation<br>RSA - behavior |
| --- | --- | --- |
| *Task set complexity* |  |  |
| Left IFG/IFJ | Z = -3.005* | r = 0.515* |
| *Response set complexity* |  |  |
| M1 / PM / SMA /<br>IPS | Z = -3.712* | r = 0.565* |
| Left PHC | Z = -3.712* | r = 0.558* |
| *Target category* |  |  |
| Left fusiform<br>gyrus/ precuneus /<br>IFG/IFJ | Z = -3.712* | r = 0.543* |
| Right MTG/TPJ | Z = -4.419* | r = 0.495* |
| Right IFG | Z = -3.712* | r = 0.533* |
| Medial SFG | Z = -2.652* | r = 0.482* |

*Note:* The asterisks indicate significance at $P < .05$ on the Wilcoxon paired-sample signed-rank test (middle column) or Pearson correlation coefficient (left column). In the last case, multiple comparisons were controlled with an FWE criterion. Abbreviations stand for inferior frontal gyrus (IFG), inferior frontal junction (IFJ), primary motor cortex (M1), premotor cortex (PM) supplementary motor area (SMA), parahippocampal cortex (PHC), middle temporal gyrus (MTG), temporoparietal junction (TPJ) and superior frontal gyrus (SFG).

### *MVPA* results

We finally aimed to explore the effect of reward directly on decoding accuracies, employing MVPA (Haxby, Connolly, & Guntupalli, 2014), as it has been previously reported during rule encoding in a classic, repetitive task-switching setting (Etzel et al., 2016). We discriminated between the two conditions of each instruction-related variable (i.e., one among faces and food-related trials, other for single versus sequential response requirements, and a last one for within versus across-dimension integration instructions) separately for rewarded and non-rewarded trials. We trained and tested our classifiers across the whole brain using searchlight and obtained, as a result, an accuracy map for each motivation condition and variable. Nonetheless, while classification was above chance in different brain regions for the three variables, we did not detect any differences in accuracies between rewarded and non-rewarded trials, as no cluster survived at the group-level the *t-*test assessing above zero differences between the two motivation conditions.

## Discussion

In the present study, we aimed to characterize the representational space for novel instructions during their proactive preparation. We assessed whether variables linked to proactive control organized encoding activity patterns and whether this structure was affected by reward expectations. Our results portrayed a complex landscape, where different organizational principles governed instruction encoding in FP cortices and lower-level perceptual and motor areas.

The left IFG/IFJ reflected the most complex and overarching representational structure, with activity patterns structured by dimension integration, response complexity and target category. Robust evidence supports the role of the IFJ in task-set reconfiguration (Brass, Derrfuss, Forstmann, & Cramon, 2005) in practiced (e.g. Woolgar, Hampshire, Thompson, & Duncan, 2011) and novel contexts (e.g. González-García et al., 2016; Muhle-Karbe et al., 2017), orchestrating neural dynamics during attentional selection (e.g. Baldauf & Desimone, 2014). This region seems to be involved in task-set maintenance (Sakai, 2008), selecting task-relevant information represented in perceptual regions (Cole, Reynolds, et al., 2013; Miller & Cohen, 2001). The current study advances our knowledge about the structure underlying *how* information is coded during novel instruction encoding, and stresses the diversity of task parameters that orchestrate task encoding in the IFG/IFJ. Such a complex, multidimensional representational space (Rigotti et al., 2013) could be key to support the richness and flexibility of human behavior in novel environments. This perspective qualifies recent research, based on MVPA, that highlights the compositionality characterizing representations held in the IFG (Cole, Laurent, et al., 2013; Deraeve, Vassena, & Alexander, 2019; Reverberi, Görgen, & Haynes, 2012), by which complex tasks are coded by combining their simpler constituent elements.

The IPS also encoded novel rules proactively, but now according to response complexity. While this is quite consistent with previous studies linking the parietal cortex to action preparation, it is worth noticing the distinction found in our data between parietal and prefrontal regions, a finding further confirmed with a more sensitive ROI analysis. Dimension integration, the variable manipulated to appeal to a higher-level task goal representation, had an effect only on LPFC, while the IPS was linked to the more specific response-set complexity (De Baene & Brass, 2014; Rubinstein et al., 2001). The frequent coativation of IFG/IFJ and IPS in demanding paradigms (Duncan, 2010) had complicated the identification of their separate contributions. The differential pattern we observed is highly relevant to disentangle their proactive role. Interestingly, the emerging picture portraits the IFG/IFJ and the IPS collaborating during novel task representation, with the former maintaining overarching representations of all relevant variables, and the latter activating the relevant stimulus-response contingencies (see also Muhle-Karbe et al., 2014). The use of RSA in our paradigm provides a deeper understanding of this process, emphasizing that the proposed two-stage preparatory mechanism also guides task-set encoding in FP cortices. In this sense, variables key for abstract goal or specific S-R settings become relevant differentially depending on the region.

Additional medial and lateral frontal cortices also participate in the FP network and are frequently recruited during task preparation (Duncan, 2010). Consequently, we also examined instruction coding in these MD regions. Our findings highlighted other LPFC areas reflecting target category (both the RLPFC and MFG) and dimension integration (MFG). The overall pattern of results obtained both with whole-brain and with ROI approaches reflects high heterogeneity within the FP network in general, and in the LPFC in particular, in terms of the attributes structuring task-set representation. In contrast, we did not obtain evidence supporting proactive task-set encoding in the ACC/preSMA and the aIfO regions. This finding fits with the subdivision of the FP network into two differentiated components: one anchored in the LPFC and IPS, and a second one composed by the ACC and the aIfO (Dosenbach et al., 2007; Palenciano et al., 2018). In line with our results, anticipatory task coding has been predominantly found in regions from the former rather than in the latter (Crittenden, Mitchell, & Duncan, 2016). Ultimately, the variability found within the FP control network during proactive novel task setting (Palenciano et al., 2018), with different processes and representational formats being combined, could be key to maximize flexibility.

Fronto-parietal cortices were not the sole brain regions encoding novel instruction parameters. Activity in fusiform gyri was organized according to target category, whereas patterns in somatomotor cortices reflected response complexity. While these regions are not associated *per se* with proactive control, their involvement reflects that their representational geometry is tuned in an anticipatory fashion by relevant task parameters conveyed by instructions. It is important to stress that all the results discussed were locked to instruction encoding, where no target stimuli had been presented, neither any specific motor response could have been prepared. These findings suggest that FP areas exert a bias in posterior cortices, according to the content of instructions. Supporting this, increments of mean activity (Esterman & Yantis, 2010) and target-specific information encoding (e.g. Stokes, Thompson, Nobre, & Duncan, 2009) have been reported in perceptual and motor regions during preparation. Importantly, these changes have been linked to boosts in functional connectivity between the FP and posterior cortices (González-García et al., 2016; Sakai & Passingham, 2006). In direct relation to our findings, a recent study showed that the representational organization in regions along the visual pathway is dynamically adapted to task demands (Nastase et al., 2017). Our current results add to these findings by showing that representational space tuning could be a mechanism of preparatory bias, which could reflect predictive coding principles where iterative loops of feedback and feedforward communication shape cognition (Friston, 2005).

Crucially, the structure of information encoded by all these regions was sensitive to trial-wise motivational states. Surprisingly, reward expectation diminished the dissimilarities between the representations of the instructions although preserving the organizational scheme found in each area. Based on recent findings of increased task decodability (Etzel et al., 2016), we had hypothesized that reward would either polarize the representational structure or increase the representational distances overall. Results were, however, in the opposite direction, even when our reward manipulation was successful at boosting performance and also increased activity in control and reward-related regions (Parro et al., 2017). Most importantly, decreases in dissimilarities were also robustly correlated with behavioral improvements. Taking into account that additional analysis employing MVPA and using data from the implementation stage corroborated these results, their implication must be thoughtfully considered. One possibility is that the decrease in dissimilarities is generated by a general boost of reward in signal-to-noise ratio. Although our results persisted after normalizing data across trials, a reward-related reduction of multivariate noise pattern could still be possible, and it could benefit task coding in the absence of the hypothesized RSA results. However, the MVPA did not reveal improved task classification accuracy in the rewarded condition, and thus this interpretation remains uncertain. Alternatively, motivation could have influenced task coding in ways that our searchlight procedure was not sensitive to. That would be the case if reward affected the spatial distribution of information: as ROIs were defined by size-fixed searchlight spheres, and were equal in rewarded and non-rewarded conditions, an effect like that would remain shadowed. Finally, the task complexity could also be key. In less demanding situations such as repetitive task switching (Etzel et al., 2016), reward could directly sharpen task encoding representations. In novel environments, however, motivation could exert a more general effect at the process level -instead of at the representational one. It could increase the efficiency of task reconfiguration (Braem & Egner, 2018), as indexed by the improvements in behavior, while the specific rule representations would remain equally structured. Nonetheless, more research is needed to properly characterize the intricate interactions among proactive control and motivation (Pessoa, 2017) in rich task environments, more akin to daily life situations.

The current study entails some limitations that constrain the scope of our findings and call for further research. On the one hand, the nature of our paradigm demanded the selection of a few instruction-organizing variables. Some other dimensions, critical for anticipatory encoding, may have been left unaddressed. Furthermore, non-linear combinations of variables could add to the organization principles governing control regions (Rigotti et al., 2013). Considering an increasing number of plausible models in more complex and/or naturalistic scenarios, together with data-driven methods such as multidimensional scaling or component analysis, will complement our results. On the other hand, our main dependent variable (fMRI hemodynamic signal) provided spatially precise, but temporal impoverished data. Temporally resolved techniques, such as electroencephalography or magnetoencephalography, could be key to unveil the temporal dynamics of the representational patterns.

Overall, our findings provide novel insights on how verbal complex novel instructions organize proactive brain activations. The emerging picture departs from pure localizationist approaches where brain regions carry fixed information about concrete cognitive processes. Rather, the different dimensions relevant for efficient instructed action shape brain activity across an extended set of areas, flexibly structuring encoding activity according to the relevant task parameters.

## Acknowledgments

This work was supported by the Spanish Ministry of Science and Innovation (PSI2016-78236-P) and the Spanish Education, Culture and Sports Ministry (FPU2014/04271 and EST16/00772 to A.F.P.). This research is part of A.F.P.’s activities for the Psychology Graduate Program of the University of Granada. We are grateful to Srikanth Padmala for his valuable help during the planning and implementation of the different fMRI data analysis employed in the current experiment.

